# Reactive astrocyte-driven epileptogenesis is induced by microglia initially activated following status epilepticus

**DOI:** 10.1101/806398

**Authors:** Fumikazu Sano, Eiji Shigetomi, Youichi Shinozaki, Haruka Tsuzukiyama, Kozo Saito, Katsuhiko Mikoshiba, Kanji Sugita, Masao Aihara, Schuichi Koizumi

## Abstract

Extensive activation of glial cells during a latent period has been well documented in various animal models of epilepsy; however, it remains unknown whether such glial activation is capable of promoting epileptogenesis. Here, we show that temporally distinct activation profiles of microglia and astrocytes collaboratively contribute to epileptogenesis in a drug-induced status epilepticus model. We found that reactive microglia appear first, followed by reactive astrocytes and increased susceptibility to seizures. Pharmacological intervention against microglial activation reduces astrogliosis, aberrant astrocyte Ca^2+^ signaling, and seizure susceptibility. Reactive astrocytes exhibit larger Ca^2+^ signals mediated by IP_3_R2, whereas deletion of this type of Ca^2+^ signaling reduces seizure susceptibility after status epilepticus. Together, our findings indicate that the sequential activation of glial cells constitutes a cause of epileptogenesis after status epilepticus.

## Introduction

Epileptogenesis; i.e., the process leading to epilepsy, is a common sequel of brain insults such as brain injury, cerebrovascular disease, or status epilepticus (SE) [1, 2] Such brain insults are typically followed by a latent period, during which the brain undergoes a cascade of morphological and functional changes over month to years prior to the onset of chronic epilepsy [3, 4]. Extensive activation of glial cells, including microglia and astrocytes, has been well documented during this latent period in various animal models of epilepsy [5–7]. Although the association of pathology with reactive glial cells is widely recognized, it is unclear whether such microglial and astrocytic activation constitutes primary causes of epilepsy or rather represents the results of repeated seizures. Moreover, the potential for these reactive glial cells to comprise candidates for epileptogenesis raises the further mechanistic question regarding whether activated glial cells might contribute to epileptogenesis independently or collaboratively.

In chemoconvulsant-induced epilepsy models, microglia are activated and produce pro-inflammatory mediators immediately following seizure onset [8]. Activated microglia can decrease the seizure threshold in animal models by releasing pro-inflammatory molecules with neuromodulatory properties [9]. Notably, the extent of microglial activation correlates with the seizure frequency in human drug-resistant epilepsy [10]. Alternatively, such microglial activation may not persist chronically. For example, pro-inflammatory molecules are detectable in microglia following a seizure but the expression diminishes after several hours [11]. Furthermore, although the activation of microglia is well characterized, it is unclear whether these activated microglia affect developing epileptogenic processes directly or through the modulation of other cells, such as subsequent astrocytic activation.

Reactive astrogliosis is also one of the most common pathological features in epilepsy and other brain insults [12, 13]. Although reactive astrogliosis is considered the consequence of repetitive seizures, some evidence that reactive astrocytes may be responsible for repetitive seizures is available. In the epileptic brain, reactive astrocytes exhibit physiological and molecular changes, such as reduced inward rectifying K^+^ current [14], changes in transporters [15], or release of gliotransmitters [16] that may underlie neuronal hyperexcitability [17]. Although astrocytes do not exhibit prominent electrical excitability as observed in neurons, they are able to dynamically regulate calcium using internal stores [18, 19]. Calcium transients in astrocytes are thought to modulate the release of a number of gliotransmitters that could influence synaptic function, synapse formation [20–22], and neural circuit excitability [23]. In particular, several previous studies showed that astrocyte calcium activity could contribute to excitotoxic neuronal death through glutamate release following SE [24, 25]. However, the functional changes including Ca^2+^ signaling of reactive astrocytes after SE and their causal roles in epileptogenesis remain largely uncertain.

To evaluate the role of inter-glial communication between different types of glial cells in the process of epileptogenesis, we assessed the spatiotemporal dynamics of glial activation following SE. Using cell-type specific manipulation, we show that relative alterations of both microglia and astrocytes play causal roles in epileptogenesis. Moreover, reactive glia are temporally distinct and collaboratively contribute to epileptogenesis. Reactive microglia appear first and induce reactive astrocytes in the hippocampus after SE. These reactive astrocytes present larger IP_3_R2-mediated Ca^2+^ signals, which are essential for induction of the increased seizure susceptibility after SE. We clearly demonstrate that inhibition of microglial activation reduces astrogliosis, aberrant astrocytic Ca^2+^ signaling, and seizure susceptibility. We therefore conclude that the sequential activation of glial cells; i.e., the initial activation of microglia followed by astrocytic activation, is a cause of epileptogenesis after SE.

## Results

### Astrocytic activation follows microglial activation after SE

To determine the contributions of glial cells to epileptogenesis, we used the pilocarpine model of epilepsy in mice, a model known to be highly isomorphic with human temporal lobe epilepsy [26, 27]. Repeated low doses of pilocarpine (100 mg kg^−1^) were injected intraperitoneally (i.p.) until the onset of SE (Fig 1A). This ramping protocol has been shown to reduce mortality after SE [28, 29]. To investigate how glial cell activation affects the epileptogenic process, we first examined the spatiotemporal pattern of microglial and astrocytic activation in the hippocampus following SE. We initially assessed microglial and astrocytic activation with immunohistochemistry using cell-type-specific activation markers at 1, 3, 7, and 28 days after SE (Fig 1B and 1D). The area of Iba1-positive microglia was significantly increased in CA1 from 1 to 7 days after SE, which was followed by an increase in the area of GFAP-positive astrocytes in CA1 from 7 to 28 days after SE (Fig 1C and 1E).

**Fig 1.**
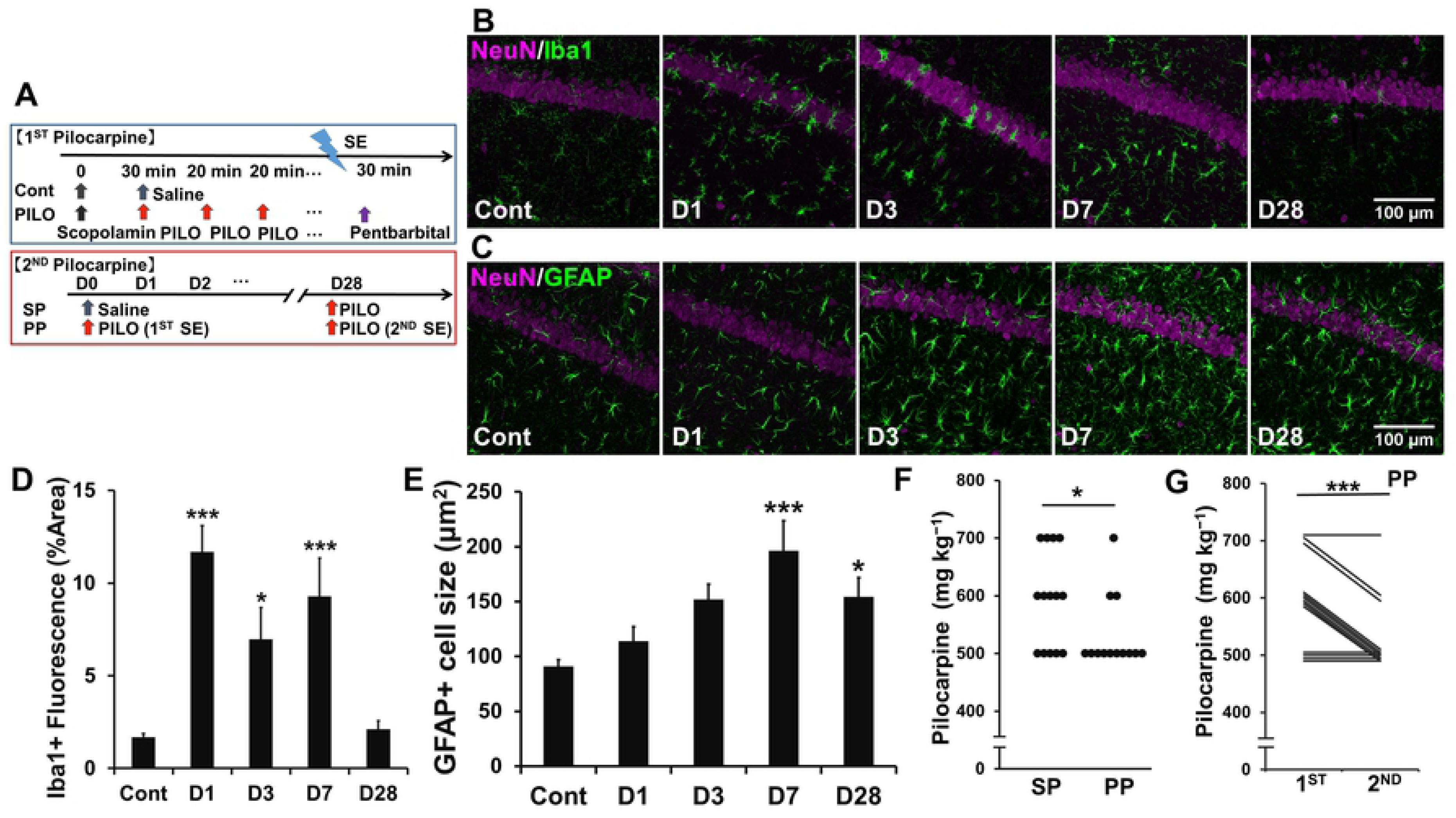
Astrogliosis is observed following microglial activation after SE. (A) As shown in the experimental protocols, mice were administered pilocarpine to achieve stage 5 seizures. The second SE was induced using the same protocol 4 weeks after the first SE. SP (PP) indicates that mice were injected with saline (pilocarpine) at 8 weeks of age followed by an injection of pilocarpine at 12 weeks of age. (B and C) Representative microphotographs showing the spatiotemporal characteristics of Iba-1 (B) or GFAP (C) expression in CA1 after SE. Fifteen images were captured per z-stack image (0.5 μm step). Cont, control; D, day. (D and E) Quantification of the temporal profile of Iba-1 positive microglia (D) or GFAP positive astrocytes (E) after SE (n = 5 mice (D); n = 5, 5, 5, 5, 7 mice, (E), \**P* < 0.05, \*\**P* < 0.01 vs. control, one-way ANOVA (*P* < 0.001) with Dunnett’s test). (F) Dot plots showing dose of pilocarpine required for the induction of the second SE (n = 14, 13 mice, \**P* < 0.05, Mann–Whitney U-test). (G) Scatter plot showing dose of pilocarpine required for the induction of the first (at 8 weeks of age) and second (at 12 weeks of age) SE in the PP group (n = 13 mice, \*\**P* < 0.01, Wilcoxon signed-rank test). Values represent the means ± SEM.

To examine whether the first SE increased seizure susceptibility, the second SE was induced 4 weeks after the first SE. A lower dose of pilocarpine was required for the induction of the second SE in mice with prior exposure to pilocarpine-induced SE at 8 weeks of age (PP) compared to those without such exposure (SP) (Fig 1F). In addition, a lower dose of pilocarpine was required for the induction of the second SE compared to the first SE (Fig 1G). These data indicated that the first SE increased seizure susceptibility at 4 weeks after the first SE. A comparison with the results in Fig 1 suggested that the temporal pattern of astrocyte activation, rather than that of microglia, correlates well with the increase of seizure susceptibility.

### Ca^2+^ hyperactivity via IP_3_R2 in reactive astrocytes after SE

To examine the SE-induced functional changes in astrocytes, Ca^2+^ imaging was performed from hippocampal slices prepared from wild-type (WT) and Glast-CreERT2::flx-GCaMP3 mice [30, 31]. Astrocytes displayed significantly larger and more frequent Ca^2+^ signals approximately 4 weeks after SE (S1 Movie). To test whether hyperactivity of astrocytes is influenced by neuronal hyperactivity, we blocked neuronal transmission by topically applying the voltage-gated sodium channel blocker tetrodotoxin (TTX; 1 µM). TTX did not affect the amplitude of astrocytic Ca^2+^ signals (Fig 2A, 2D and 2E) (S2 Movie).

**Fig 2.**
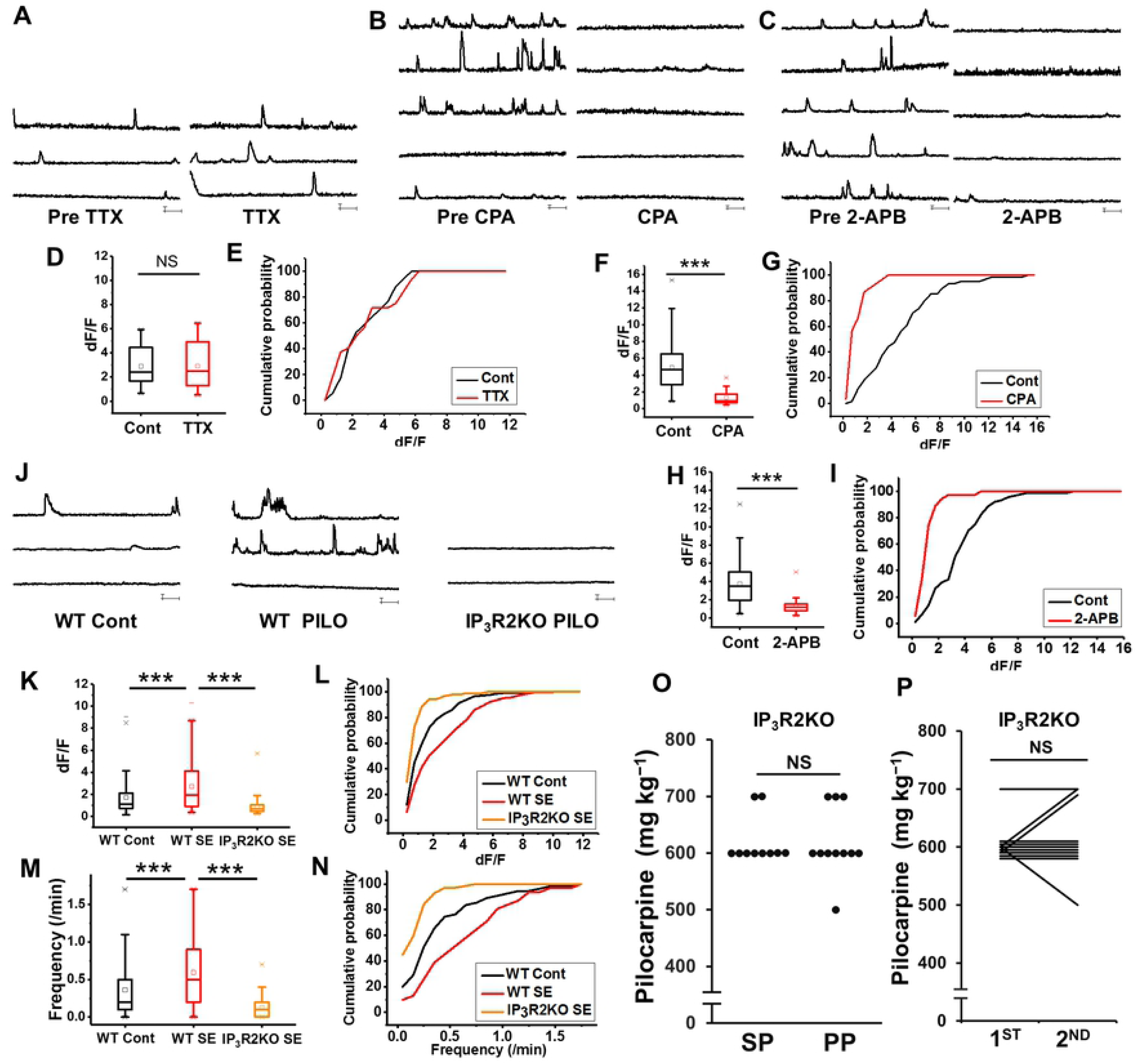
Reactive astrocytes exhibit IP_3_R2-mediated Ca^2+^ hyperactivity, which is essential for epileptogenesis. (A-C) Ca^2+^ dynamics of astrocytes approximately 4 weeks after SE in the CA1 stratum radiatum region in Glast-CreERT2::flx-GCaMP3 mice before and after TTX (1 μM) (A), CPA (20 μM) (B), and 2-APB (100 μM) (C) application. (D-I) Box plots showing amplitudes of Ca^2+^ signals before and after TTX (1 μM) (D), CPA (20 μM) (F), and 2-APB (100 μM) (H) application. (n = 10, 13, 14 cells/2 mice, \*\*\**P* < 0.001, unpaired t-test). Cont, control. Cumulative probability plots showing amplitudes (dF/F) of Ca^2+^ signals before and after TTX (not significant (*P* > 0.05), Kolmogorov–Smirnov test) (E), CPA (*P* < 0.001, Kolmogorov– Smirnov test) (G), and 2-APB (*P* < 0.001, Kolmogorov–Smirnov test) (**I**) application. (J) Astrocytic Ca^2+^ dynamics by Fluo4 in the CA1 stratum radiatum region in WT control, WT after SE, and IP_3_R2KO mice after SE. (K-N) Box plots showing Ca^2+^ signal amplitudes (dF/F) (K) and frequency (M) (n = 57, 32, 85 cells/2, 2, 3 mice, \*\*\**P* < 0.001, unpaired t-test). Cumulative probability plots showing Ca^2+^ signal amplitudes (dF/F) (L) and frequency (N) (*P* < 0.001, Kolmogorov–Smirnov test). (O) Dot plots showing dose of pilocarpine required for the induction of the second SE in IP_3_R2KO mice. SP (PP) indicates mice were injected with saline (pilocarpine) at 8 weeks of age followed by an injection of pilocarpine at 12 weeks of age. (n = 10 mice, N.S., not significant (*P* > 0.05), Mann–Whitney U-test). (P) Scatter plot showing dose of pilocarpine required for the induction of the first (at 8 weeks of age) and second (at 12 weeks of age) SE in the PP group regarding IP3R2KO mice (n = 10 mice, N.S., not significant (*P* > 0.05), Wilcoxon signed-rank test). Note: The first pilocarpine did not affect the dose required for the second SE in IP_3_R2KO, see Fig 1G.

To elucidate the molecular mechanisms involved in astrocytic Ca^2+^ hyperactivity, we applied cyclopiazonic acid (CPA; 20 μM) to deplete intracellular calcium stores. CPA significantly reduced the amplitude of astrocytic Ca^2+^ signals after SE (Fig 2B, 2F and G) (S2 Movie). Then, we applied the membrane-permeable IP_3_ receptor antagonist 2-aminoethoxydiphenyl borate (2-APB; 100 μM). 2-APB also significantly reduced the amplitude of astrocytic Ca^2+^ signals after SE (Fig 2C, 2H and 2I) (S4 Movie). To confirm that astrocytic Ca^2+^ hyperactivity is completely dependent on the IP_3_ receptor, we performed Ca^2+^ imaging in IP_3_R2 knockout (KO) mice [32]. The amplitude of astrocytic Ca^2+^ signals after SE was significantly decreased in IP_3_R2KO mice compared with that in WT (Fig 2J, 2K, 2L, 2M and 2N). The frequency of astrocytic Ca^2+^ signals after SE was also significantly decreased in IP_3_R2KO mice (Fig 2M and 2N) (S5 Movie). These results suggested that astrocytic Ca^2+^ hyperactivity after SE should be dependent on IP_3_R2-mediated Ca^2+^ release from internal stores.

### IP_3_R2KO mice exhibit rescue of the increased seizure susceptibility

To clarify the role of astrocytic Ca^2+^ hyperactivity after SE in epileptogenesis, we investigated seizure susceptibility after SE in IP_3_R2KO mice [32]. No differences in the dose of pilocarpine required for the induction of the first SE were observed between IP_3_R2KO and WT mice (Fig 1F and 2O). These data indicated that IP_3_R2-mediated Ca^2+^ signaling in astrocytes does not alter the acute responses to pilocarpine.

In IP_3_R2KO mice, the area of Iba1-positive microglia was significantly increased in CA1 at 1 day after SE, suggesting that microglial activation after SE was comparable in IP_3_R2KO and WT mice (S1 Fig). However, there was no significant change in the dose of pilocarpine required for the induction of the second SE in SP compared with PP mice (Fig 2O). There was no significant change in the dose of pilocarpine required for the induction of the first and second SE in IP_3_R2KO mice (Fig 2P). These results suggested that IP_3_R2-mediated astrocytic Ca^2+^ hyperactivity is essential for the induction of the increased seizure susceptibility after SE.

### Microglia inhibition reduces activated astrocyte morphology

Our data indicated temporal differences between activation of microglia and astrocytes; i.e., earlier and later onset after SE, respectively. To reveal features of the activated microglia after SE, we investigated the changes in mRNA levels of pro-inflammatory cytokines that are relevant to microglial activation by quantitative reverse transcription-polymerase chain reaction (RT-PCR) (Fig 3A, 3B and 3C). SE increased *Tnf* and *Il1b* mRNA in the hippocampus at 1 day after SE (Fig 3B and 3C). To explore the microglia-triggered astrocyte activation, we investigated microglial functional changes after SE. Among several molecules tested, we found that *Tnf* and *Il1b* mRNAs were also significantly upregulated in the isolated hippocampal microglia at 1 day after SE (Fig 3A).

**Fig 3.**
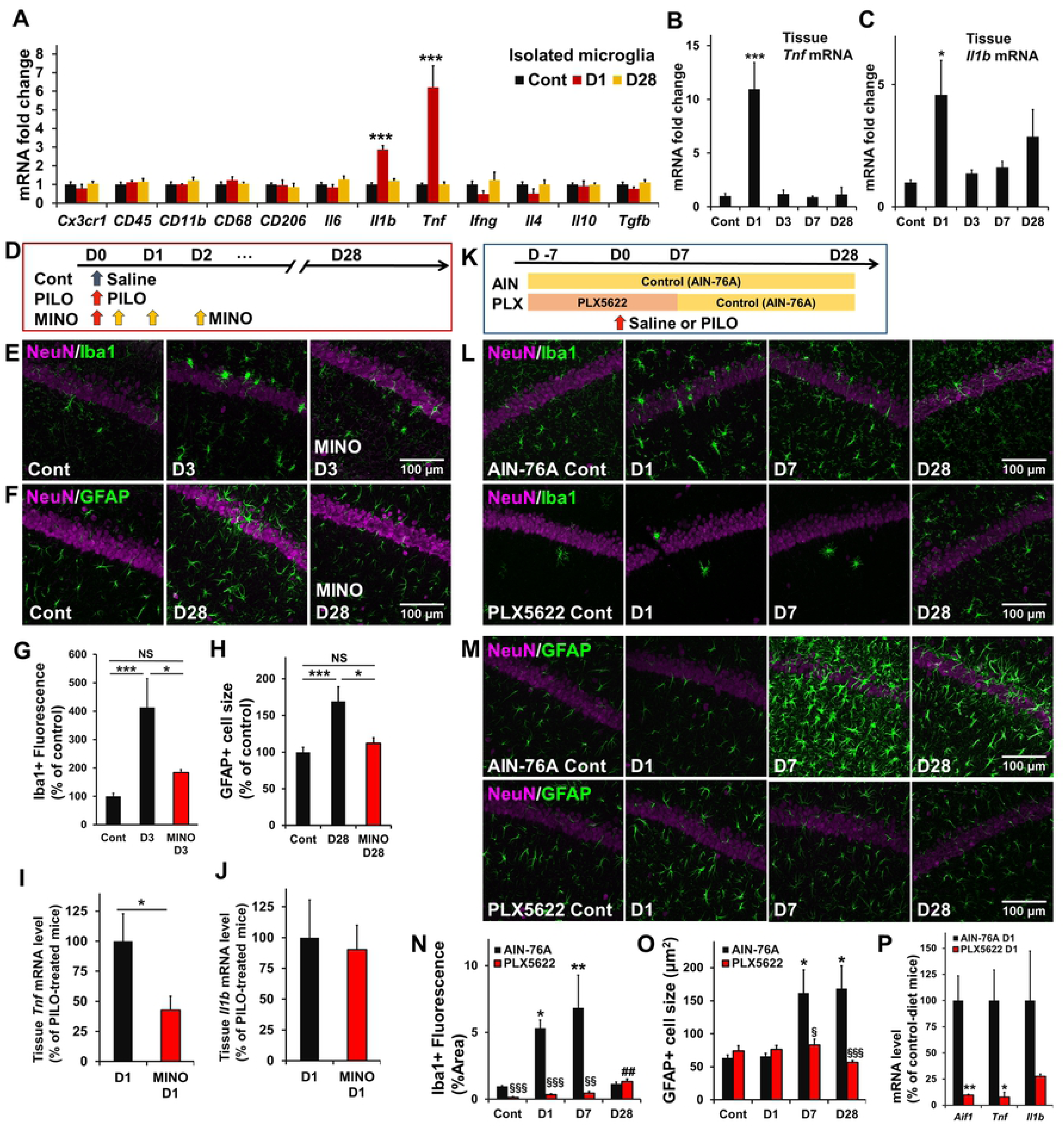
Microglia inhibition with minocycline and depletion with CSF1R antagonist (PLX5622) reduces astrogliosis. (A) Microfluidic quantitative RT-PCR analysis of mRNA in total RNA extracted from hippocampal microglia after SE. Relative ratios of *Gapdh-*normalized mRNA to the corresponding control (Cont) are shown (n = 3 samples/9 mice, \*\*\**P* < 0.001 vs. control, one-way ANOVA (*P* < 0.01, *P* < 0.001) with Dunnett’s test). (B and C) Quantitative RT-PCR analysis of *Tnf* and *ll1b* mRNA in total hippocampal RNA after SE. Relative ratios of *Gapdh*-normalized mRNA to the corresponding control are shown (n = 5 mice, \**P* < 0.05, \*\*\**P* < 0.001 vs. control, one-way ANOVA (*P* < 0.001, *P* < 0.05) with Dunnett’s test). (D) Experimental scheme for minocycline post-treatment-mediated microglia inhibition. (E-H) Representative microphotographs showing the spatiotemporal characteristics of Iba-1 (E) and GFAP (F) expression and quantification of Iba-1 positive microglia (G) and GFAP positive astrocytes (H) in CA1 with or without minocycline post-treatment after SE. Fifteen images were collected per z-stack image (0.5 μm step). (n = 5 mice, N.S., not significant (*P* > 0.05), \**P* < 0.05, \*\*\**P* < 0.001, one-way ANOVA (*P* < 0.01) with Bonferroni test). (I and J) Quantitative RT-PCR analysis as in (B and C) with or without minocycline post-treatment. (n = 5 mice, N.S., not significant (*P* > 0.05), \**P* < 0.05, unpaired t-test). (K) Experimental scheme for PLX5622-mediated microglia depletion. (L-O) Representative microphotographs showing the spatiotemporal characteristics of Iba-1 (L) and GFAP (M) expression and quantification of Iba-1 positive microglia (N) and GFAP positive astrocytes (O) in CA1 with or without PLX5622 after SE. Fifteen images were captured per z-stack image (0.5 μm step). (n = 5 mice, \**P* < 0.05, \*\**P* < 0.01 vs. control of AIN-76A (control diet), ^#*#*^*P* < 0.01 vs. control of PLX5622, ^§^*P* < 0.05, ^§§^*P* < 0.01, ^§§§^*P* < 0.001 vs. AIN-76A (corresponding day), one-way ANOVA (*P* < 0.01) with Dunnett’s test and unpaired t-test). (P) Quantitative RT-PCR analysis as in (B and C) with or without PLX5622. (n = 5 mice, \**P* < 0.05, \*\**P* < 0.01, unpaired t-test).

To clarify whether microglial activation is required for astrogliosis, we investigated the effect of post-treatment with the inhibitor, minocycline (Fig 3D) [33–35]. To confirm the efficacy of minocycline in this protocol, microglial activation was assessed by immunohistochemistry and quantitative RT-PCR. Minocycline post-treatment prevented the increase in the area of Iba1-positive cells in CA1 at 3 days after the first SE (Fig 3E and 3G) along with an increase in *Tnf* but not *Il1b* mRNA in the hippocampus at 1 day after the first SE (Fig 3I and 3J). Notably, microglia inhibition with minocycline post-treatment prevented the increase in the area of GFAP-positive cells in CA1 at 28 days after the first SE (Fig 3F and 3H).

To further confirm that acute microglial activation plays an important role in the morphological activation of astrocytes after SE, we applied PLX5622, a CSF1R antagonist, to deplete microglia (Fig 3K) [36–38]. PLX5622 treatment prevented the increase in the area of Iba1-positive cells in CA1 from 1 to 7 days after the first SE (Fig 3L and 3N). In addition, *Aif1* and *Tnf* mRNA levels were significantly decreased at 1 day after SE with PLX5622 treatment compared with those in the control diet group (Fig 3P). Similarly, the increased area of GFAP-positive astrocytes in CA1 from 7 to 28 days after SE in control diet (AIN-76A) mice was prevented in PLX5622 treated mice (Fig 3M and 3O). To identify the optimal timing of microglial inhibition to prevent astrogliosis, we applied PLX5622 from 3 weeks after SE (Fig 4A). This later PLX5622 treatment decreased the area of Iba1-positive cells in CA1 at 28 days after the first SE (Fig 4B and 4D) but did not prevent the increased area of GFAP-positive astrocytes (Fig 4C and 4E). These findings showed that the initial reactive microglia are required to induce morphological activation of astrocytes after SE.

**Fig 4.**
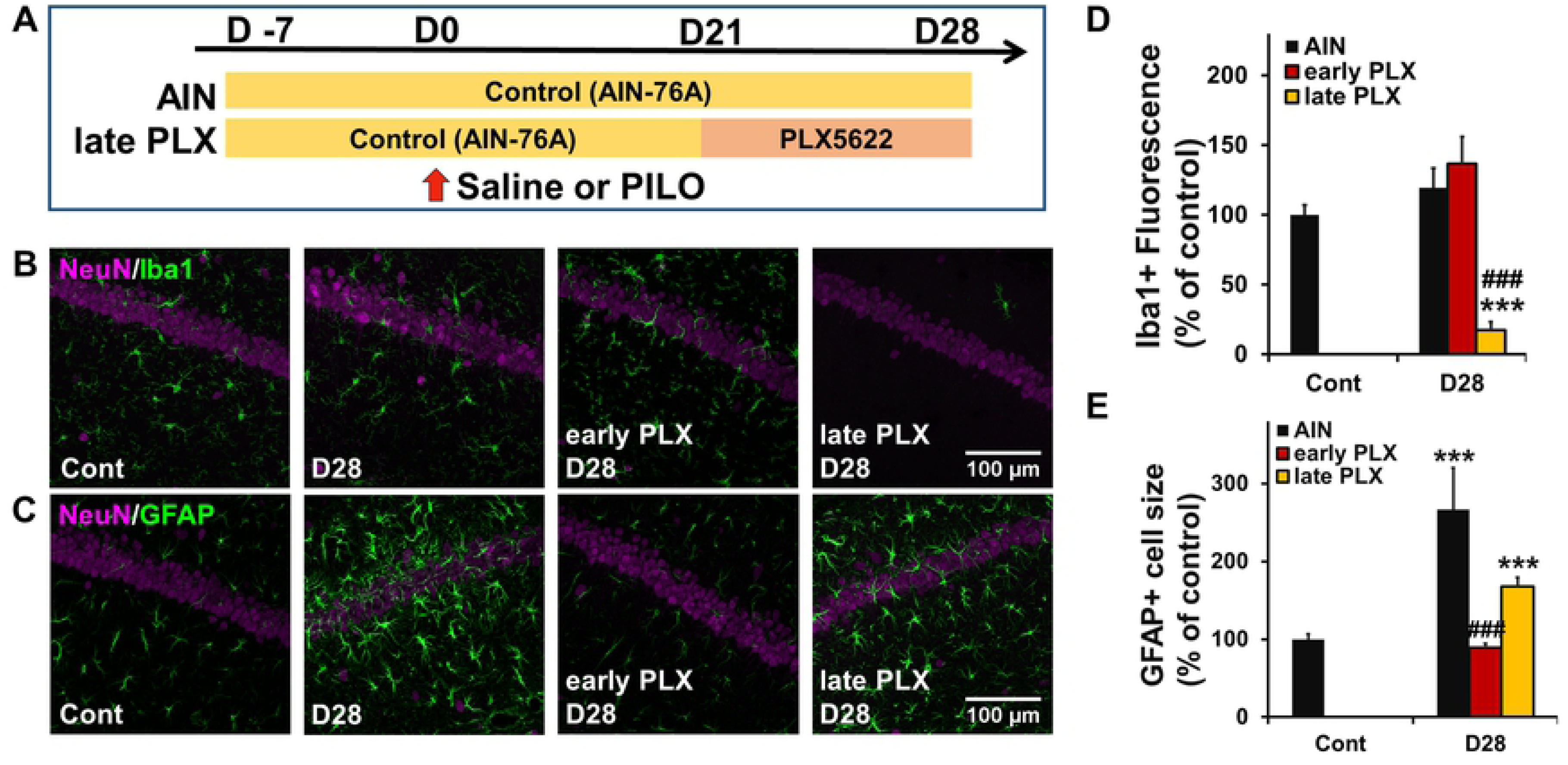
Microglia depletion with CSF1R antagonist (PLX5622) at late phase after SE does not reduce astrogliosis and increased seizure susceptibility. (A) Experimental scheme for microglia depletion with PLX5622 at the late phase after SE. (B and C) Representative microphotographs showing the spatiotemporal feature of Iba-1 (B) and GFAP (C) expression in CA1 with or without PLX5622 after SE. Fifteen images were collected per z-stack image (0.5 μm step). Cont, control; D, day. (D and E) Quantification of the temporal profile of Iba-1 positive microglia (D) and GFAP positive astrocytes (E) after SE (n = 5 mice, \*\*\**P* < 0.01 vs. control, unpaired t-test, ^##*#*^*P* < 0.01 vs. AIN-76A (corresponding day), one-way ANOVA (*P* < 0.001) with Dunnett’s test). Values represent the means ± SEM.

### Microglia inhibition reduces astrocytic Ca^2+^ hyperactivity

We then investigated whether microglial activation is required for astrocytic Ca^2+^ hyperactivity after SE. We also used a pharmacological approach to inhibit the early microglial activation after SE. Microglia inhibition with minocycline reduced the larger and frequent Ca^2+^ signals of astrocytes (S1 Movie) (Fig 5A, 5B, 5C, 5D and 5E). Similarly, the amplitude and frequency of fluo-4AM-labeled astrocytic Ca^2+^ signaling after SE were significantly increased in control diet (AIN-76A) mice (Fig 5F, 5H, 5I, 5J and 5K) (S6 Movie). Conversely, the larger and frequent Ca^2+^ signals after SE were significantly reduced by the PLX5622 treatment (Fig 5G, 5L, 5M, 5N and 5O) (S7 Movie). These results indicated that acute microglial activation is essential for the changes of astrocytic Ca^2+^ activity after SE.

**Fig 5.**
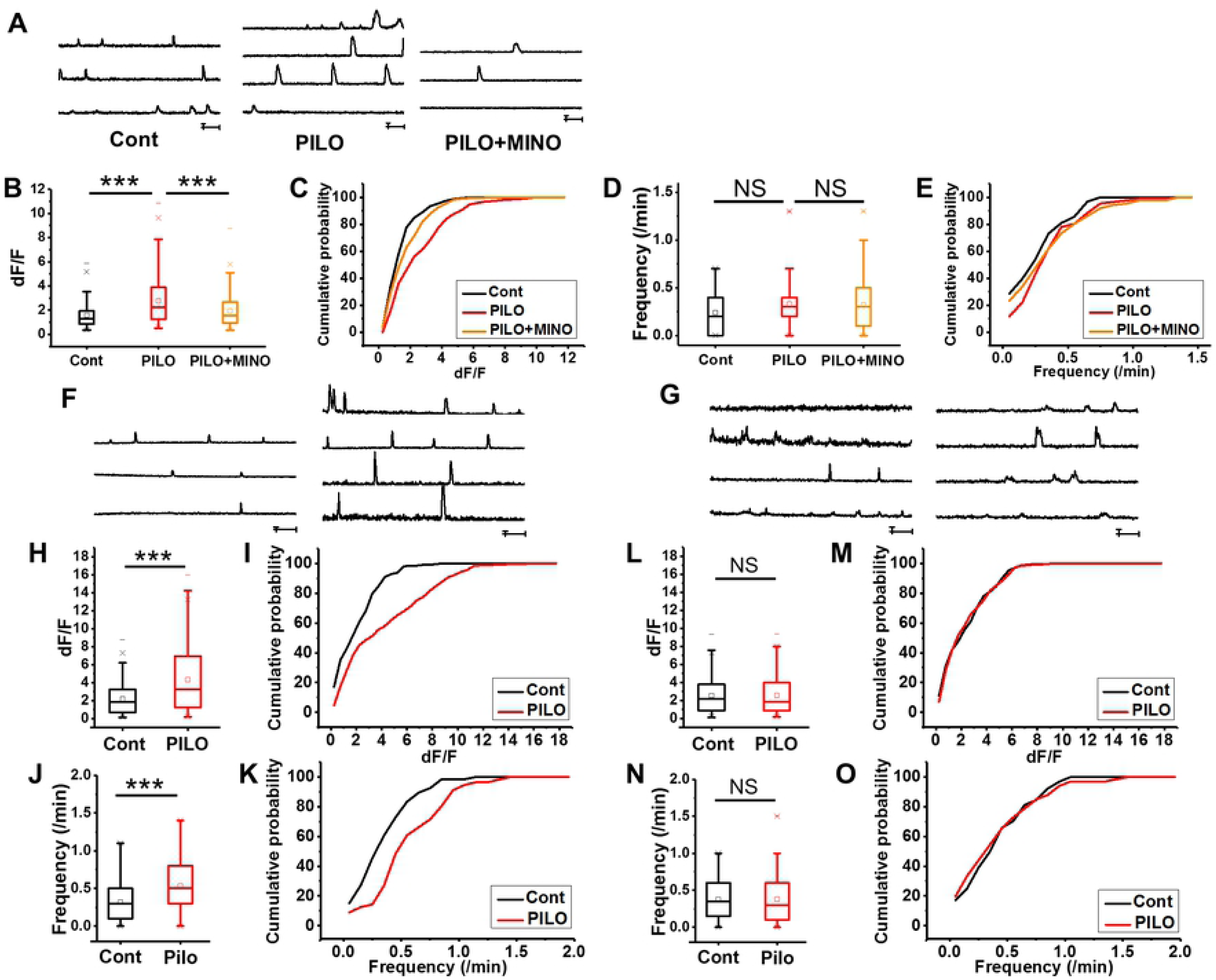
Microglia inhibition with minocycline or CSF1R antagonist (PLX5622) reduces the increased astrocytic Ca^2+^ hyperactivity following SE. (A) Ca^2+^ dynamics of astrocytes approximately 4 weeks after SE in the CA1 stratum radiatum region in Glast-CreERT2::flx-GCaMP3 mice with or without minocycline treatment. (B-E) Box plots showing Ca^2+^ signal amplitude (dF/F) (B) and frequency (D) (n = 74, 92, 93 cells/3 mice, N.S., not significant (*P* > 0.05), \*\*\**P* < 0.001, unpaired t-test). Cumulative probability plots showing Ca^2+^ signal amplitude (dF/F) (*P* < 0.001, Kolmogorov–Smirnov test) (C) and frequency (not significant (*P* > 0.05), Kolmogorov–Smirnov test) (E). (F and G) Ca^2+^ dynamics of astrocytes approximately 4 weeks after SE in the CA1 stratum radiatum region in Glast-CreERT2::flx-GCaMP3 mice with (G) or without (F) PLX5622 treatment. (H-K) Box plots showing Ca^2+^ signal amplitude (dF/F) (H) and frequency (J) in the AIN-76A (control diet) group. (n = 70, 58 cells/2 mice, \*\*\**P* < 0.001, unpaired t-test). Cumulative probability plots showing Ca^2+^ signal amplitude (dF/F) (*P* < 0.001, Kolmogorov–Smirnov test) (I) and frequency (*P* < 0.001, Kolmogorov–Smirnov test) (K) in the AIN-76A (control diet) group. (L-O) Box plots showing Ca^2+^ signal amplitude (dF/F) (L) and frequency (M) in the PLX5622 group. (n = 61, 71 cells/2 mice, N.S., not significant (*P* > 0.05), unpaired t-test). Cumulative probability plots showing Ca^2+^ signal amplitude (dF/F) (not significant (*P* > 0.05), Kolmogorov–Smirnov test) (M) and frequency (not significant (*P* > 0.05), Kolmogorov–Smirnov test) (O) in the PLX5622 group.

### Microglia inhibition rescues enhanced seizure susceptibility

Finally, we tested whether microglia inhibition rescued the increased seizure susceptibility following SE. Post-treatment with minocycline following the first SE prevented the increased seizure susceptibility (Fig 6A and 6B). No difference was observed between control diet and PLX5622-treated mice in the dose of pilocarpine required for the induction of the first SE (Fig 6C), indicating that microglia inhibition does not alter the acute responses to pilocarpine. In contrast, a lower dose of pilocarpine was required for the induction of the second SE in control mice compared with that in PLX5622-treated mice (Fig 6D). Consistent with this, unlike the enhanced seizure susceptibility observed in control mice following the first SE (as indicated by the reduced dose of pilocarpine required to induce the second vs. the first SE), there was no significant change in the dose of pilocarpine required for the induction of the first or second SE in PLX5622-treated mice (Fig 6E and 6F). In contrast, a lower dose of pilocarpine was required for the induction of the second SE in later PLX5622 treatment mice, similar to that in control diet mice (Fig 6G, 6H, 6I and 6J). These data suggested that the inhibition of initial microglial activation rescues the increased seizure susceptibility.

**Fig 6.**
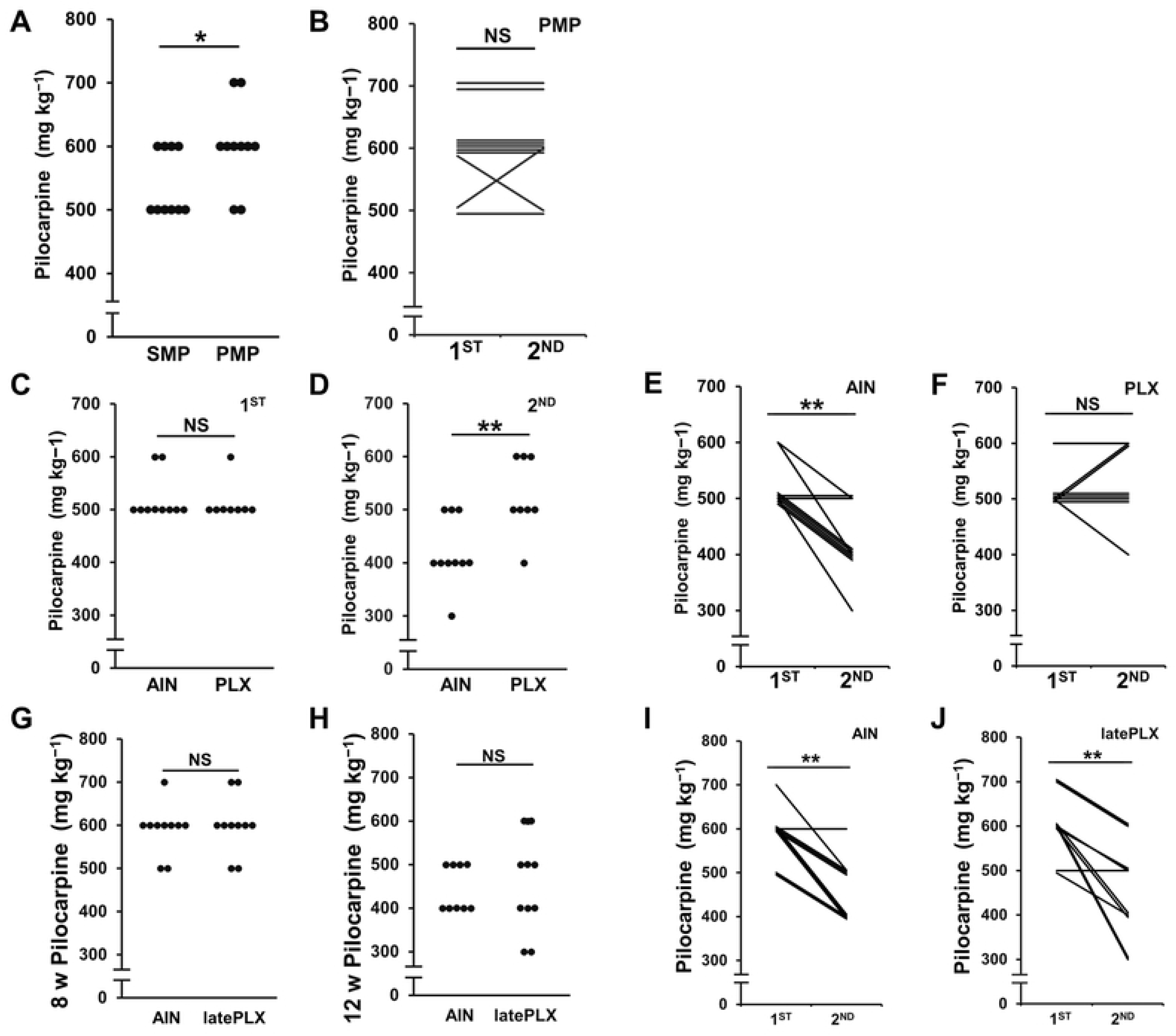
Microglia inhibition with minocycline or CSF1R antagonist (PLX5622) reduces the increased seizure susceptibility following SE. (A) Dot plots showing dose of pilocarpine required for the induction of the second SE (n = 10 mice, N.S., not significant (*P* > 0.05), \**P* < 0.05, Mann– Whitney U-test). SMP (PMP) indicates that mice were injected with saline (pilocarpine) at 8 weeks of age with minocycline post-treatment followed by an injection of pilocarpine at 12 weeks of age. (B) Scatter plot showing dose of pilocarpine required for the induction of the first (at 8 weeks of age) and second (at 12 weeks of age) SE. (n = 10 mice, \*\**P* < 0.01, Wilcoxon signed-rank test). (C and D) Dot plots showing dose of pilocarpine required for the induction of the first (C) and second (D) SE with or without PLX5622. (n = 10, 8 mice, N.S., not significant (*P* > 0.05), \*\**P* < 0.01, Mann–Whitney U-test). AIN, control diet (AIN-76A). (E and F) Scatter plot showing dose of pilocarpine required for the induction of the first (at 8 weeks of age) and second (at 12 weeks of age) SE for AIN-76A (control diet) (E) or PLX5622 (F). (n = 10, 8 mice, N.S., not significant (*P* > 0.05), \*\**P* < 0.01, Wilcoxon signed-rank test). (G and H) Dot plots showing dose of pilocarpine required for the induction of the first (G) and second (H) SE with or without late PLX5622 treatment. (n = 10 mice, N.S., not significant (*P* > 0.05), Mann–Whitney U-test). (I and J) Scatter plot showing dose of pilocarpine required for the induction of the first (at 8 weeks of age) and second (at 12 weeks of age) SE AIN-76A (control diet) (I) or PLX5622 (J). (n = 10 mice, \*\**P* < 0.01, Wilcoxon signed-rank test).

## Discussion

Here, we demonstrate that SE induces sequential activation of glial cells; i.e., the initial activation of microglia, followed by astrocytic activation, which is essential for seizure susceptibility or epileptogenesis. The main findings in the present study are as follows. (1) Microglia are activated and pro-inflammatory cytokines of microglia are increased immediately after SE. (2) Reactive astrocytes, which exhibit larger IP_3_R2-mediated Ca^2+^ signals, appear following microglial activation after SE. (3) Genetic deletion of IP_3_R2 rescues both the aberrant Ca^2+^ signals in astrocytes and the increased seizure susceptibility. (4) Pharmacological inhibition of microglial activation or deletion of microglia reduces astrogliosis along with aberrant Ca^2+^ signals of astrocytes, and rescues the increased seizure susceptibility. These findings indicate that initially activated microglia are responsible for the subsequent induction of epileptogenic reactive astrocytes in vivo.

Microglial and astrocytic activation is a common feature of various central nervous system (CNS) disorders including epilepsy [39–42]. However, the pathological significance and spatiotemporal pattern of microglial and astrocytic activation in the epileptogenic process have not been carefully addressed. Microglial response to SE occurs immediately, with reactive microglia playing both detrimental and beneficial roles during acute seizures [43]. Although activated microglia exhibit a neuroprotective role via the P2Y12 receptor in the acute phase, they exert proconvulsive effects through the production of pro-inflammatory cytokines such as IL-1β [11], TNF [44], and IL-6 [45, 46]. However, such increase of purinergic receptors and pro-inflammatory cytokines after SE may be transient [11], and it is unknown how this transient microglial activation including pro-inflammatory cytokines causes long-term epileptic potential. Here, we found that inhibiting microglia at the acute phase (0 to 7 days after SE) but not the late phase (21 to 28 days after SE) reduced susceptibility to the second SE, suggesting that activated microglia trigger the epileptogenic process including astrocytic activation, but do not exert a direct proconvulsive effect on the later phase after SE.

In the present study, we demonstrate that astrocytic activation develops slowly starting 7 days after SE, is long lasting, and still observed when mice show increased seizure susceptibility. Astrogliosis is thought to contribute to the pathophysiology of epilepsy [47–49]. However, the role of astrogliosis in epileptogenesis is largely unknown. In particular, it is important to determine whether activated astrocytes play a proconvulsive or anticonvulsive role in the epileptic brain. It has been proposed that astrocytic Ca^2+^ signaling contributes to the induction of epileptic seizures and neuronal cell loss by seizures [27,28,50,51]. In this study, we observed larger Ca^2+^ signals in the somatic regions of astrocytes in the latent phase of epileptogenesis. Analysis of the Ca^2+^ signals in astrocytes suggests that these Ca^2+^ signals are mediated by IP_3_R2. Notably, we found that genetic deletion of IP_3_R2 is sufficient to rescue the increased seizure susceptibility and reduce astrogliosis. Our study thus suggests that IP_3_R2-mediated Ca^2+^ signaling in reactive astrocytes plays a proconvulsive role in the epileptic brain and can contribute to epileptogenesis.

Astrocytic Ca^2+^ signals may contribute to epileptogenesis through several mechanisms. Astrocytes impact neural circuit excitability directly by releasing “gliotransmitters”, such as glutamate [23,52,53]. Astrocytes also increase neuronal excitability by forming new circuits through the release of synaptogenic molecules [22, 54]. However, the functional consequences of these changes in the context of epileptogenesis remain to be determined. As Ca^2+^ serves as a ubiquitous intracellular signal in the regulation of numerous cellular processes including exocytosis, proliferation, and gene expression, it is also likely to regulate many processes in the induction/maintenance of reactive astrocytes [55, 56]. Although it is difficult to exclude the inherent influence on the neural circuit resulting from the deletion of IP_3_R2, we demonstrate that SE induces neither an increase in Ca^2+^ excitation in astrocytes nor proconvulsive effects in IP_3_R2KO mice, suggesting that enhanced Ca^2+^ signals in astrocytes are likely responsible for epileptogenesis.

In animal models of epilepsy, reactive astrocytes undergo extensive physiological changes involving not only Ca^2+^ signaling but also ion and neurotransmitter homeostasis along with intracellular and extracellular water content, which can cause neuronal hyperexcitability [57–60]. The relative importance of such functional changes of astrocytes to epileptogenesis will be investigated in future studies. Recently, it has been reported that activated microglia can induce neurotoxic reactive astrocytes (i.e., A1 astrocytes), which release unidentified neurotoxic factors [37, 61]. Thus, whether astrogliosis after SE results in a similar phenotype to A1 astrocytes and whether IP_3_-mediated Ca^2+^ signals contribute to the induction of neurotoxic phenotype [56] represent relevant issues to be addressed in future investigations. Although whether the astrocytes induced by activated microglia are in a primarily neurotoxic or neuroprotective state remains largely unknown, our data suggest that the reactive astrocytes induced by activated microglia after SE exert proconvulsive effects in the epileptic brain.

In this study, we also demonstrate that pro-inflammatory cytokines of microglia are increased prior to astrocytic activation, suggesting the importance of microglial activation as an initial process of epileptogenesis. Pharmacological inhibition and depletion of microglia significantly blocked the activation of astrocytes and decreased the seizure threshold after SE. Our findings identify that activated microglia likely promote epileptogenesis by inducing the proconvulsive phenotype of astrocytes. Although it has been recognized that microglial activation occurs before reactive astrogliosis in various CNS diseases [62—64], little was known prior to the present study regarding how microglial-astrocytic interactions contribute to the pathophysiology of epilepsy. For example, several previous studies using chemoconvulsant-induced epilepsy models have shown that activated microglia were present immediately after SE and that functional changes occurred, such as upregulation of pro-inflammatory cytokines [8,65,66], purinergic receptors [39], and phagocytosis [40].

Previous reports also revealed that microglia modulate astrocyte activation via various molecules, especially pro-inflammatory cytokines [67, 68]. Consistent with this, we found that TNF and IL-1β are significantly upregulated in hippocampal microglia at 1 day after SE. Conversely, microglia inhibition by minocycline prevents the increased mRNA of TNF in the hippocampus at 1 day after the first SE along with subsequent reactive astrogliosis, suggesting a potential role of pro-inflammatory cytokines from microglia in reactive astrogliosis after SE. As the effect of minocycline may not be restricted to microglia, we depleted microglia using a CSF-1 receptor antagonist and found similar results, suggesting that microglial activation occurs through cytokine release. Thus, despite the potential problem of specificity owing to the use of pharmacological inhibition of microglia, we clearly show that initial activation of microglia and microglia-derived proinflammatory cytokines likely underlie the subsequent astrogliosis-mediated epileptogenesis. Nevertheless, because the molecular mechanisms underlying the activation of astrocytes triggered by activated microglia have not been fully clarified, other chemical mediators such as ATP may also contribute to activate microglia-mediated astrogliosis [69]. Further investigations using more specific interventions are required to elucidate the precise molecular mechanisms underlying the interaction between microglia and astrocytes.

In summary, our findings identify a sequence of glial activation in the hippocampus that contributes to the epileptogenic process. In this process, microglial activation is identified as a crucial event to induce reactive astrocytes. In turn, astrocytic Ca^2+^ activation mediated by IP_3_R2 was essential for the induction of epileptogenesis. Our findings add to the emerging view that reactive astrocytes triggered by microglia have a central role in the pathogenesis of epilepsy and, given the limited progress of neuron-centered epilepsy research over the past several years, suggest reactive astrocytes as promising new targets for the development of alternative and more specific antiepileptic drugs.

## Materials and Methods

### Animals

All studies used male C57BL/6J mice (SLC Japan, Shizuoka, Japan). IP_3_R2KO mice on a C57BL/6 background were available from a previous study [32]; their generation and maintenance have been previously described in detail. Glast-CreERT2::flx-GCaMP3 mice on a C57BL/6 background were also available from a previous study [30, 31]; their generation and maintenance have been previously described in detail. In the present study, we performed immunohistochemistry and confirmed that GCaMP3 was co-localized with GFAP, an astrocyte marker, but not with Iba1 or NeuN (S2 Fig and S1 Table). Overall, Ca^2+^ signals detected by GCaMP3 were mainly detected from astrocytes.

Mice were housed on a 12 h light (6 am)/dark (6 pm) cycle with ad libitum access to water and rodent chow. The animals were allowed to adapt to laboratory conditions for at least 1 week before starting the experiments. All experimental procedures were performed in accordance with the “Guiding Principles in the Care and Use of Animals in the Field of Physiologic Sciences” published by the Physiologic Society of Japan and with the previous approval of the Animal Care Committee of Yamanashi University (Chuo, Yamanashi, Japan).

### Animal treatments

The first SE was induced in 8-week-old male mice by the administration of pilocarpine and the second SE was induced 4 weeks after the first SE. A low dose of 100 mg kg^−1^ pilocarpine (Wako, 161-07201) per injection was administered i.p. every 20 min until the onset of Racine scale stage 5 seizures. Scoporamin methyl bromide (1 mg kg^−1^, i.p., Wako, 198-07971) was administered 30 min prior to pilocarpine injection to reduce its peripheral effects [28, 29]. Seizures were terminated with pentobarbital (20 mg kg^−1^, i.p., Kyoritu Seiyaku) when mice experienced stage 5 seizures for 30 min. Behavior of pilocarpine-treated mice was observed for 1 h after SE.

To establish whether minocycline inhibits acute seizure-induced microglial activation, mice were administered i.p. with saline or minocycline (25 mg kg^−1^) 1 h after pilocarpine-SE induction and for the following two consecutive days [33–35]. Microglia were also depleted from mice by treatment with the CSF1R antagonist, PLX5622 (Plexxikon), formulated in AIN-76A rodent chow (Research Diets). Mice were treated with PLX5622 (1200 mg kg^−1^ Chow) or a matched control diet (AIN-76A) for seven days before SE and for the following seven consecutive days [36–38].

### Immunohistochemistry

The mice were deeply anesthetized with pentobarbital and perfused transcardially with phosphate buffered saline (PBS), followed by 4% (w/v) paraformaldehyde in PBS. The brains were removed, postfixed overnight, then cryoprotected with 30% (w/v) sucrose in PBS for two days. The brains were frozen and coronal sections (20 μm) were cut using a cryostat (Leica CM1100). Slices were washed with PBS three times and treated with 0.1% Triton-X100/10% NGS for 1 h to block nonspecific binding. The sections were incubated for two days at 4 °C with the following primary antibodies: monoclonal rat anti-GFAP (1:2000; Thermo Fisher Scientific, 13-0300), monoclonal mouse anti-NeuN (1:500; Millipore, MAB377), polyclonal rabbit anti-Iba1 (1:1000; Wako, 019-19741), chicken anti-GFP antibody (1:1000, Thermo Fisher Scientific, A10262), and rabbit anti-NeuN (1:1,000; Millipore, MABN140). The sections were washed three times with PBS and then incubated for 2 h at room temperature with secondary antibodies: Alexa 488- or Alexa 546-conjugated goat anti-mouse/rat/rabbit or chicken IgGs (1:500; Invitrogen, A11029/Thermo Fisher Scientific, A-11081/Invitrogen, A11035/Thermo Fisher Scientific, A11039). After washing slices with PBS three times, they were mounted with Vectashield Mounting Medium (Vector Laboratories). Fluorescence images were obtained using a confocal laser microscope system (FV-1000; Olympus) or Keyence fluorescence microscope (BZX-700).

### Standard quantitative RT-PCR

Total RNA was isolated and purified from tissues using the RNeasy Lipid Tissue Mini Kit (Qiagen) according to the manufacturer’s instructions. RT-PCR amplifications were performed using the One Step PrimeScript RT-PCR Kit (TaKaRa Bio). RT-PCR amplifications and real-time detection were performed using an Applied Biosystems 7500 Real-Time PCR System. The thermocycling parameters were as follows: 5 min at 42 °C for reverse transcription, 10 s at 95 °C for inactivation of the RT enzyme, and 40 cycles of denaturation (5 s at 95 °C) and annealing/extension (34 s at 60 °C). All primer probe sets and reagents were purchased from Applied Biosystems: rodent *Gapdh* (4308313), mouse *Tnf* (Mm00443260_g1), mouse *Il1b* (Mm00434228_m1).

### Dissociated cell suspensions from adult mouse brain

Three 8-week old male mice were perfused with PBS after anesthesia to eliminate serum vesicles and hippocampi were dissected to comprise one sample. Tissue dissociation was performed using the gentleMACS dissociator and the Adult Brain Dissociation Kit (Miltenyi Biotec) according to the manufacturer’s protocol. Briefly, brain tissue was minced and digested with a proprietary enzyme solution on the gentleMACS dissociator adult brain program. The cells were then incubated with anti-mouse CD11b-coated microbeads (Miltenyi Biotec) for 10 min at 4 °C. The cell-bead mix was then washed to remove unbound beads. Prior to antibody labeling, nonspecific binding to the Fc receptor was blocked using the FcR Blocking Reagent (Miltenyi Biotec). Cells were suspended in PBS with 0.5% bovine serum albumin and the cell suspension was loaded onto an LS Column (Miltenyi Biotec), which was placed in the magnetic field of a QuadroMACS™ Separator (Miltenyi Biotec). The magnetically labeled CD11b positive cells were retained within the column and eluted as the positively selected cell fraction after removing the column from the magnet.

### Microfluidic quantitative RT-PCR

Total RNA was extracted from dissociated cells using the RNeasy Lipid Tissue Mini Kit (Qiagen) and cDNA synthesis performed using the PrimeScript RT-PCR Kit (Perfect Real Time) (TaKaRa Bio). For pre-amplification, up to 100 qPCR assays (primer/probe sets in 20x stock concentration) were pooled and diluted to a 0.2x concentration. For microfluidic qPCR, 1.25 μl of each cDNA sample was pre-amplified using 1 μl of TaqMan pre-amplification master mix (PN 100-5580, Fluidigm), 1.25 μl of the primer pool, and 1.5 μl of water. Pre-amplification was performed using a 2 min 95 °C denaturation step and 14 cycles of 15 s at 95 °C and 4 min at 60 °C. Microfluidic quantitative RT-PCR reactions were performed using the 96x96 chips and included 2–3 technical replicates for each combination of sample and assay. For sample mixtures, 2.7 μl pre-amplification product was combined with 0.3 μl of 20x GE Sample Loading Reagent (85000746, Fluidigm) and 3 μl of 2x PCR master mix (4324020, Thermo Fisher Scientific), of which 5 μl of was loaded into sample wells. For assay mixtures, equal volumes of TaqMan assay and 2x Assay Loading Reagent (PN85000736, Fluidigm) were combined, and 5 μl of the resulting mixture was loaded into multiple assay wells. RT-PCR amplifications and real-time detection were performed using the BioMarkHD Real-Time PCR System (Fluidigm). Data from Fluidigm runs were manually checked for reaction quality prior to analysis, and Ct values for each gene target were normalized to Ct values for housekeeping genes. All primer probe sets and reagents were purchased from Integrated DNA Technologies: rodent *Gapdh* (Mm.PT.39a.1), mouse *Tnf* (Mm.PT.58.12575861), mouse *Il1b* (Mm.PT.58.41616450), mouse *Cx3cr1* (Mm.PT.58.17555544), mouse *CD45* (Mm.PT.58.7583849), mouse *CD11b* (Mm.PT.58.14195622), mouse *CD68* (Mm.PT.58.32698807), mouse *CD206* (Mm.PT.58.42560062), mouse *Il6* (Mm.PT.58.10005566), mouse *Ifng* (Mm.PT.58.41769240), mouse *Il4* (Mm.PT.58.32703659), mouse *Il10* (Mm.PT.58.13531087), and mouse *Tgfb* (Mm.PT.58.11254750).

### Preparation of brain slices and Ca^2+^ imaging

The methods used have been described previously [56, 70]. Briefly, 8-week-old male mice were anesthetized with pentobarbital (100 mg kg^−1^, i.p.). Cold cutting ACSF, composed of 92 mM NaCl, 2.5 mM KCl, 1.2 mM NaH_2_PO_4_, 30 mM NaHCO_3_, 20 mM HEPES, 25 mM D-glucose, 5 mM sodium ascorbate, 2 mM thiourea, 3 mM sodium pyruvate, 10 mM MgCl_2_, and 0.5 mM CaCl_2_ saturated with 95% O_2_–5% CO_2_, was perfused transcardially. Coronal slices of the hippocampus (300 µm) were cut using a vibrating microtome (Pro7, Dosaka) in cutting ACSF. Slices were incubated at 34 °C for 10 min in recovery ACSF, composed of 93 mM N-methyl-D-glucamine, 93 mM HCl, 2.5 mM KCl, 1.2 mM NaH_2_PO_4_, 30 mM NaHCO_3_, 20 mM HEPES, 25 mM D-glucose, 5 mM sodium ascorbate, 2 mM thiourea, 3 mM sodium pyruvate, 10 mM MgCl_2_, and 0.5 mM CaCl_2_ saturated with 95% O_2_–5% CO_2_, and subsequently stored in ACSF comprising 124 mM NaCl, 2.5 mM KCl, 1.2 mM NaH_2_PO_4_, 24 mM NaHCO_3_, 5 mM HEPES, 12.5 mM D-glucose, 5 mM sodium ascorbate, 2 mM thiourea, 3 mM sodium pyruvate, 2 mM MgCl_2_, and 2 mM CaCl_2_ saturated with 95% O_2_ and 5% CO_2_ at room temperature. After 1 h of recovery, slices were submerged in ACSF at approximately 32 °C. Slices were imaged using an Olympus Fluoview FVMPE-RS two-photon laser scanning microscope equipped with a Maitai HP DS-OL laser (Spectra-Physics). We used a 920 nm laser and 510 nm high-pass emission filter. Astrocytes were selected from the CA1 stratum radiatum region and were typically 30–50 μm from the slice surface. Images were gathered using a 25× water immersion lens with a numerical aperture of 1.05.

For Fluo-4AM measurements, we dropped 2.5 μl Fluo-4AM (2 mM) onto the hippocampal slices followed by incubation in ACSF for 60 min, then transferred the slices to dye-free ACSF for at least 30 min prior to experimentation. The final concentration of Fluo4-AM was 5 μM with 0.02% Pluronic F–127. Astrocytes were selected from the CA1 stratum radiatum region and were typically 30–50 μm from the slice surface. TTX (1 μM), 2-APB (100 μM), and CPA (20 μM) were solubilized in ACSF. Baseline astrocytic activity was recorded prior to drug application. Subsequently, drugs were applied onto the slice for 10 min and astrocytic activity was recorded for 10 min.

### Image analysis

Images were acquired using inverted confocal laser-scanning systems (Olympus FV-1000) at 40× magnification with a 1.30 numerical aperture objective lens. Information regarding z-stack images is described in the figure legends. Astrocytes were selected from the CA1 stratum radiatum region and imaged based on GFAP immunostaining. Microglia were imaged based on Iba1 immunostaining at the CA1 stratum radiatum region. Subsequent images were processed and quantified using ImageJ (US National Institutes of Health; NIH). For the quantitative analysis of the area containing Iba1 positive microglia, we randomly chose three fields per mouse. Images were converted to gray scale and the quantification threshold was set constantly for all specimens within each experimental group. The percentage of Iba1-positive area was calculated by dividing the area of Iba1-positive region by the total area of the region of interest. For the quantitative analysis of GFAP positive cell size, we randomly chose one field per mouse. Images were converted to gray scale and the quantification threshold was set constantly for all specimens within each experimental group. To quantify the GFAP positive cell size, particles were analyzed based on GFAP immunoreactivity and we chose the three largest GFAP positive cells per field.

The methods used for Ca^2+^ imaging data analysis have been described previously [56, 70]. Briefly, imaging data were analyzed using ImageJ. We selected regions of interest from somatic regions of astrocytes by visual examination of the time lapse image. Using these regions of interest, raw fluorescence intensity values (F) were taken from the original videos and converted to delta F/F (dF/F) in Originlab (Origin Lab Corp.). We analyzed Ca^2+^ signals when their dF/F values were greater than 0.2. We analyzed Ca^2+^ signals and their amplitude (dF/F) and duration (full width at half maximum) using the Originlab “peak analysis” function.

### Statistical analysis

All statistical analyses were performed using SPSS version 19.0 (SPSS Inc.) software. Data are presented as the means ± SEM. Most data were analyzed using one-way ANOVA followed by Dunnett’s multiple post hoc test for comparing more than three samples, and two-sample unpaired *t*-tests. *P* values <0.05 were considered as statistically significant.

## Acknowledgements

We thank Dr. K. Takanashi, Mr. R. Komatsu, Mrs. Y. Fukasawa, Mrs. M. Tachibana, Mrs. Y. Koseki, and Mrs. Y. Hoshino (Univ. Yamanashi) for technical assistance, and all members of the Koizumi Laboratory for critical discussion.

## Supporting Information

**S1 Fig** Initial microglial activation is observed after SE in IP3R2KO mice.

**S2 Fig** Immunohistochemical analysis of GCaMP expression in the hippocampus in Glast-CreERT2::Flx-GCaMP3 mice.

**S1 Table** Cell-specific markers in GCaMP3-expressing cells in the hippocampus of Glast-CreERT2::Flx-GCaMP3 mice (tamoxifen i.p. at P7).

**S1 Movie** Ca^2+^ dynamics of astrocytes in the CA1 stratum radiatum region in Glast-CreERT2::flx-GCaMP3 control mice and approximately 4 weeks after SE with or without minocycline treatment.

**S2 Movie** Ca^2+^ dynamics of astrocytes approximately 4 weeks after SE in the CA1 stratum radiatum region in Glast-CreERT2::flx-GCaMP3 mice before and after TTX application.

**S3 Movie** Ca^2+^ dynamics of astrocytes approximately 4 weeks after SE in the CA1 stratum radiatum region in Glast-CreERT2::flx-GCaMP3 mice before and after CPA application.

**S4 Movie** Ca^2+^ dynamics of astrocytes approximately 4 weeks after SE in the CA1 stratum radiatum region in Glast-CreERT2::flx-GCaMP3 mice before and after 2-APB application.

**S5 Movie** Astrocytic Ca^2+^ dynamics as revealed by Fluo4 in the CA1 stratum radiatum region in WT control, WT after SE, and IP_3_R2KO mice after SE.

**S6 Movie** Ca^2+^ dynamics of astrocytes approximately 4 weeks after SE in the CA1 stratum radiatum region in Glast-CreERT2::flx-GCaMP3 mice.

**S7 Movie** Ca^2+^ dynamics of astrocytes approximately 4 weeks after SE in the CA1 stratum radiatum region in Glast-CreERT2::flx-GCaMP3 mice with or without PLX5622 treatment.

## References

1. Engel J Jr. Mesial temporal lobe epilepsy: what have we learned? Neuroscientist. 2001;7(4): 340–52.

2. Herman ST. Epilepsy after brain insult: targeting epileptogenesis. Neurology. 2002;59(9 Suppl 5): S21-26.

3. French JA, Williamson PD, Thadani VM, Darcey TM, Mattson RH, Spencer SS, et al. Characteristics of medial temporal lobe epilepsy: I. Results of history and physical examination. Ann Neurol. 1993;34(6): 774–80.

4. Rakhade SN, Jensen FE. Epileptogenesis in the immature brain: emerging mechanisms. Nat Rev Neurol. 2009;5(7): 380–391.

5. Binder DK, Steinhauser C. Functional changes in astroglial cells in epilepsy. Glia. 2006;54(5): 358–368.

6. Seifert G, Steinhauser C. Neuron-astrocyte signaling and epilepsy. Exp Neurol. 2013;244: 4–10.

7. Shapiro LA, Wang L, Ribak CE. Rapid astrocyte and microglial activation following pilocarpine-induced seizures in rats. Epilepsia. 2008;49 Suppl 2: 33–41.

8. Benson MJ, Manzanero S, Borges K. Complex alterations in microglial M1/M2 markers during the development of epilepsy in two mouse models. Epilepsia. 2015;56(6): 895–905.

9. Vezzani A, French J, Bartfai T, Baram TZ. The role of inflammation in epilepsy. Nat Rev Neurol. 2011;7(1): 31–40.

10. Boer K, Spliet WG, van Rijen PC, Redeker S, Troost D, Aronica E. Evidence of activated microglia in focal cortical dysplasia. J Neuroimmunol. 2006;173(1-2): 188–195.

11. Vezzani A, Moneta D, Richichi C, Aliprandi M, Burrows SJ, Ravizza T, et al. Functional role of inflammatory cytokines and antiinflammatory molecules in seizures and epileptogenesis. Epilepsia. 2002;43 Suppl 5: 30–35.

12. Cendes F, Sakamoto AC, Spreafico R, Bingaman W, Becker AJ. Epilepsies associated with hippocampal sclerosis. Acta Neuropathol. 2014;128(1): 21–37.

13. Morizawa YM, Hirayama Y, Ohno N, Shibata S, Shigetomi E, Sui Y, et al. Author Correction: Reactive astrocytes function as phagocytes after brain ischemia via ABCA1-mediated pathway. Nat Commun. 2017;8(1): 1598.

14. Haj-Yasein NN, Jensen V, Vindedal GF, Gundersen GA, Klungland A, Ottersen OP, et al. Evidence that compromised K+ spatial buffering contributes to the epileptogenic effect of mutations in the human Kir4.1 gene (KCNJ10). Glia. 2011;59(11): 1635–1642.

15. Tanaka K, Watase K, Manabe T, Yamada K, Watanabe M, Takahashi K, et al. Epilepsy and exacerbation of brain injury in mice lacking the glutamate transporter GLT-1. Science. 1997;276(5319): 1699-1702.

16. Bezzi P, Gundersen V, Galbete JL, Seifert G, Steinhauser C, Pilati E, et al. Astrocytes contain a vesicular compartment that is competent for regulated exocytosis of glutamate. Nat Neurosci. 2004;7(6): 613–620.

17. Devinsky O, Vezzani A, Najjar S, De Lanerolle NC, Rogawski MA. Glia and epilepsy: excitability and inflammation. Trends Neurosci. 2013;36(3): 174–184.

18. Charles AC, Merrill JE, Dirksen ER, Sanderson MJ. Intercellular signaling in glial cells: calcium waves and oscillations in response to mechanical stimulation and glutamate. Neuron. 1991;6(6): 983–992.

19. Cornell-Bell AH, Finkbeiner SM, Cooper MS, Smith SJ. Glutamate induces calcium waves in cultured astrocytes: long-range glial signaling. Science. 1990;247(4941): 470-473.

20. Allen NJ, Eroglu C. Cell biology of astrocyte-synapse interactions. Neuron. 2017;96(3): 697–708.

21. Araque A, Carmignoto G, Haydon PG, Oliet SH, Robitaille R, Volterra A. Gliotransmitters travel in time and space. Neuron. 2014;81(4): 728–739.

22. Kim SK, Hayashi H, Ishikawa T, Shibata K, Shigetomi E, Shinozaki Y, et al. Cortical astrocytes rewire somatosensory cortical circuits for peripheral neuropathic pain. J Clin Invest. 2016;126(5): 1983–1997.

23. Halassa MM, Fellin T, Haydon PG. The tripartite synapse: roles for gliotransmission in health and disease. Trends Mol Medi. 2007;13(2): 54–63.

24. Ding S, Fellin T, Zhu Y, Lee SY, Auberson YP, Meaney DF, et al. Enhanced astrocytic Ca2+ signals contribute to neuronal excitotoxicity after status epilepticus. J Neurosci. 2007;27(40): 10674–10684.

25. Tian GF, Azmi H, Takano T, Xu Q, Peng W, Lin J, et al. An astrocytic basis of epilepsy. Nat Med. 2005;11(9): 973–981.

26. Curia G, Longo D, Biagini G, Jones RS, Avoli M. The pilocarpine model of temporal lobe epilepsy. J Neurosci Methods. 2008;172(2): 143–157.

27. Levesque M, Avoli M, Bernard C. Animal models of temporal lobe epilepsy following systemic chemoconvulsant administration. J Neurosc Methods. 2016;260: 45–52.

28. Clasadonte J, Dong J, Hines DJ, Haydon PG. Astrocyte control of synaptic NMDA receptors contributes to the progressive development of temporal lobe epilepsy. Proc Natl Acad Sci U S A. 2013;110(43): 17540–17545.

29. Groticke I, Hoffmann K, Loscher W. Behavioral alterations in a mouse model of temporal lobe epilepsy induced by intrahippocampal injection of kainate. Exp Neurol. 2008;213(1): 71–83.

30. Mori T, Tanaka K, Buffo A, Wurst W, Kuhn R, Gotz M. Inducible gene deletion in astroglia and radial glia--a valuable tool for functional and lineage analysis. Glia. 2006;54(1): 21–34.

31. Zariwala HA, Borghuis BG, Hoogland TM, Madisen L, Tian L, De Zeeuw CI, et al. A Cre-dependent GCaMP3 reporter mouse for neuronal imaging in vivo. J Neurosci. 2012;32(9): 3131–3141.

32. Futatsugi A, Nakamura T, Yamada MK, Ebisui E, Nakamura K, Uchida K, et al. IP3 receptor types 2 and 3 mediate exocrine secretion underlying energy metabolism. Science. 2005;309(5744): 2232-2234.

33. Abraham J, Fox PD, Condello C, Bartolini A, Koh S. Minocycline attenuates microglia activation and blocks the long-term epileptogenic effects of early-life seizures. Neurobiol Dis. 2012;46(2): 425–430.

34. Hirayama Y, Ikeda-Matsuo Y, Notomi S, Enaida H, Kinouchi H, Koizumi S. Astrocyte-mediated ischemic tolerance. J Neurosci. 2015;35(9): 3794–3805.

35. Matsuda T, Murao N, Katano Y, Juliandi B, Kohyama J, Akira S, et al. TLR9 signalling in microglia attenuates seizure-induced aberrant neurogenesis in the adult hippocampus. Nat Commun. 2015;6: 6514.

36. Dagher NN, Najafi AR, Kayala KM, Elmore MR, White TE, Medeiros R, et al. Colony-stimulating factor 1 receptor inhibition prevents microglial plaque association and improves cognition in 3xTg-AD mice. J Neuroinflamm. 2015;12: 139.

37. Shinozaki Y, Shibata K, Yoshida K, Shigetomi E, Gachet C, Ikenaka K, et al. Transformation of astrocytes to a neuroprotective phenotype by microglia via P2Y1 receptor downregulation. Cell Rep. 2017;19(6): 1151–1164.

38. Valdearcos M, Robblee MM, Benjamin DI, Nomura DK, Xu AW, Koliwad SK. Microglia dictate the impact of saturated fat consumption on hypothalamic inflammation and neuronal function. Cell Rep. 2014;9(6): 2124–2138.

39. Avignone E, Ulmann L, Levavasseur F, Rassendren F, Audinat E. Status epilepticus induces a particular microglial activation state characterized by enhanced purinergic signaling. J Neurosci. 2008;28(37): 9133–9144.

40. Koizumi S, Shigemoto-Mogami Y, Nasu-Tada K, Shinozaki Y, Ohsawa K, Tsuda M, et al. UDP acting at P2Y6 receptors is a mediator of microglial phagocytosis. Nature. 2007;446(7139): 1091-1095.

41. Pekny M, Nilsson M. Astrocyte activation and reactive gliosis. Glia. 2005;50(4): 427–434.

42. Sofroniew MV, Vinters HV. Astrocytes: biology and pathology. Acta Neuropathol. 2010;119(1): 7–35.

43. Eyo UB, Murugan M, Wu LJ. Microglia-neuron communication in epilepsy. Glia. 2017;65(1): 5–18.

44. Stellwagen D, Beattie EC, Seo JY, Malenka RC. Differential regulation of AMPA receptor and GABA receptor trafficking by tumor necrosis factor-alpha. J Neurosci. 2005;25(12): 3219–3228.

45. Campbell IL, Abraham CR, Masliah E, Kemper P, Inglis JD, Oldstone MB, et al. Neurologic disease induced in transgenic mice by cerebral overexpression of interleukin 6. Proc Natl Acad Sci U S A. 1993;90(21): 10061–10065.

46. Samland H, Huitron-Resendiz S, Masliah E, Criado J, Henriksen SJ, Campbell IL. Profound increase in sensitivity to glutamatergic- but not cholinergic agonist-induced seizures in transgenic mice with astrocyte production of IL-6. J Neurosci Res. 2003;73(2): 176–187.

47. Coulter DA, Steinhauser C. Role of astrocytes in epilepsy. Cold Spring Harb Perspect Med. 2015;5(3): a022434.

48. Seifert G, Carmignoto G, Steinhauser C. Astrocyte dysfunction in epilepsy. Brain Res Rev. 2010;63(1-2): 212–221.

49. Steinhauser C, Grunnet M, Carmignoto G. Crucial role of astrocytes in temporal lobe epilepsy. Neuroscience. 2016;323: 157–169.

50. Alvarez-Ferradas C, Morales JC, Wellmann M, Nualart F, Roncagliolo M, Fuenzalida M, et al. Enhanced astroglial Ca2+ signaling increases excitatory synaptic strength in the epileptic brain. Glia. 2015;63(9): 1507–1521.

51. Gomez-Gonzalo M, Losi G, Chiavegato A, Zonta M, Cammarota M, Brondi M, et al. An excitatory loop with astrocytes contributes to drive neurons to seizure threshold. PLoS Biol. 2010;8(4): e1000352.

52. Bazargani N, Attwell D. Astrocyte calcium signaling: the third wave. Nat Neurosci. 2016;19(2): 182–189.

53. Haydon PG. GLIA: listening and talking to the synapse. Nat Rev Neurosci. 2001;2(3): 185–193.

54. Weissberg I, Wood L, Kamintsky L, Vazquez O, Milikovsky DZ, Alexander A, et al. Albumin induces excitatory synaptogenesis through astrocytic TGF-beta/ALK5 signaling in a model of acquired epilepsy following blood-brain barrier dysfunction. Neurobiol Dis. 2015;78: 115–125.

55. Berridge MJ, Bootman MD, Roderick HL. Calcium signalling: dynamics, homeostasis and remodelling. Nat Rev Mol Cell Biol. 2003;4(7): 517–529.

56. Saito K, Shigetomi E, Yasuda R, Sato R, Nakano M, Tashiro K, et al. Aberrant astrocyte Ca(2+) signals “AxCa signals” exacerbate pathological alterations in an Alexander disease model. Glia. 2018;66(5): 1053–1067.

57. Bedner P, Dupper A, Huttmann K, Muller J, Herde MK, Dublin P, et al. Astrocyte uncoupling as a cause of human temporal lobe epilepsy. Brain. 2015;138(Pt 5): 1208–1222.

58. Bordey A, Lyons SA, Hablitz JJ, Sontheimer H. Electrophysiological characteristics of reactive astrocytes in experimental cortical dysplasia. J Neurophysiol. 2001;85(4): 1719–1731.

59. Bordey A, Sontheimer H. Properties of human glial cells associated with epileptic seizure foci. Epilepsy Res. 1998;32(1-2): 286–303.

60. Steinhauser C, Seifert G, Bedner P. Astrocyte dysfunction in temporal lobe epilepsy: K+ channels and gap junction coupling. Glia. 2012;60(8): 1192–1202.

61. Liddelow SA, Guttenplan KA, Clarke LE, Bennett FC, Bohlen CJ, Schirmer L, et al. Neurotoxic reactive astrocytes are induced by activated microglia. Nature. 2017;541(7638): 481-487.

62. Dheen ST, Kaur C, Ling EA. Microglial activation and its implications in the brain diseases. Curr Med Chem. 2007;14(11): 1189–1197.

63. Kingwell K. Neurodegenerative disease: Microglia in early disease stages. Nat Rev Neurol. 2012;8(9): 475.

64. Shinozaki Y, Nomura M, Iwatsuki K, Moriyama Y, Gachet C, Koizumi S. Microglia trigger astrocyte-mediated neuroprotection via purinergic gliotransmission. Sci Repo. 2014;4: 4329.

65. Riazi K, Galic MA, Kuzmiski JB, Ho W, Sharkey KA, Pittman QJ. Microglial activation and TNFalpha production mediate altered CNS excitability following peripheral inflammation. Proc Nl Acad Sci U S A. 2008;105(44): 17151–17156.

66. Vezzani A, Conti M, De Luigi A, Ravizza T, Moneta D, Marchesi F, et al. Interleukin-1beta immunoreactivity and microglia are enhanced in the rat hippocampus by focal kainate application: functional evidence for enhancement of electrographic seizures. J Neurosci. 1999;19(12): 5054–5065.

67. John GR, Lee SC, Brosnan CF. Cytokines: powerful regulators of glial cell activation. Neuroscientist. 2003;9(1): 10–22.

68. Sofroniew MV. Molecular dissection of reactive astrogliosis and glial scar formation. Trends Neurosc. 2009;32(12): 638–647.

69. Imura Y, Morizawa Y, Komatsu R, Shibata K, Shinozaki Y, Kasai H, et al. Microglia release ATP by exocytosis. Glia. 2013;61(8): 1320–1330.

70. Shigetomi E, Hirayama YJ, Ikenaka K, Tanaka KF, Koizumi S. Role of purinergic receptor P2Y1 in spatiotemporal Ca(2+) dynamics in astrocytes. J Neurosci. 2018;38(6): 1383–1395.

